# MicroRNA miR-196a controls neural crest patterning by repressing immature neural ectoderm programs in Xenopus embryos

**DOI:** 10.1101/2025.03.05.641602

**Authors:** Alice M. Godden, Nicole Ward, Méghane Sittewelle, Marco Antonaci, Aleksandr Kotov, Anne H. Monsoro-Burq, Grant N. Wheeler

**Affiliations:** School of Biological Sciences, University of East Anglia, Norwich Research Park, Norwich, NR4 7TJ, United Kingdom; Université Paris-Saclay, Département de Biologie, Faculté des Sciences d’Orsay, Signalisation Radiobiology and Cancer, CNRS UMR 3347, INSERM U1021, Orsay F-91405, France; Institut Curie Research Division, Paris Science et Lettres Research University, Orsay F-91405; Institut Universitaire de France, Paris F-75005, France

**Keywords:** microRNAs, miR-196a, neural crest, BMP signalling, neural induction, pluripotency, Xenopus laevis and tropicalis embryos, dorsal ectoderm gene regulatory networks

## Abstract

**Highlights:** miR-196a promotes Neural Crest development:

- RNA sequencing of the developing neural border and neural crest reveals miR-196a-dependent gene programs.
- miR-196a represses early neural plate programs and reduces pluripotency gene expression.
- miR-196a may directly control neural crest gene *sox10 in vivo*.

Neural crest (NC) cells form a multipotent stem cell population specified during neurulation, which undergo an epithelial-to-mesenchymal transition (EMT) and migrate extensively in the developing embryo, to generate numerous tissues and cell types including the craniofacial skeleton, the peripheral nervous system and pigment cells. The genetic and molecular details of NC specification are governed by a complex, yet still partially understood gene regulatory network (NC-GRN). In particular, the precise function of microRNAs (miRNA) in this network remains poorly characterized. miRNAs are short non-coding 20-22 nucleotides long RNAs, which control gene expression through post-transcriptional repression. Since miRNA-196a is expressed in the developing neural and NC cells of *Xenopus laevis* embryos, we here investigated miR-196a function in the NC-GRN, by knocking-down its expression using antisense morpholinos. Depletion of miR-196a revealed major NC and craniofacial phenotypes. These defects were preceded by the perturbed expression of key neural, neural border and NC markers such as *sox2/3, zic1/3, pax3*, *sox10* and *snail2*. Using RNA sequencing of individual neural border and NC explants, we have identified a signature of genes up- and down-regulated by miR-196a and validate these with rescue experiments using a miRNA mimic. Our study identifies miR-196a as a key actor of early patterning in the dorsal ectoderm, balancing the extent of immature neural plate progenitors with NC and placode specification, while also promoting neuron differentiation within the neural plate.

## INTRODUCTION

MicroRNAs (miRNAs) are short, single-stranded, non-coding RNAs, of approximately 22 nucleotides long (Alberti and Cochella, 2017; Lee et al., 1993; Shah et al., 2017). They act mainly as repressors of gene expression by binding to the untranslated region at the 3’ end of a targeted mRNA and either promoting the stalling of the ribosome or directly promoting the degradation of the targeted mRNA. They can be found either in intronic regions of the genome, processed from the introns allowing for co-expression of the miRNA and coordinating protein (Salim et al., 2022), or as independent genes from which many miRNA precursors can be produced in one transcript and then processed individually (Thatcher et al., 2008). MirRNAs are highly conserved in evolution and are found in both plants and animals (Bartel, 2004). It was reported that 60% of protein-coding genes in humans have conserved target sites for miRNAs (Friedman et al., 2009). In particular, miRNAs play important roles in the development of invertebrates like the worm and fruit fly (Chandra et al., 2017) and of vertebrates, including chick, mouse, frog and fish (Ahmed et al., 2015; Lewis and Steel, 2010; Mok et al., 2017; Thatcher et al., 2008; Ward et al., 2018). Some miRNAs regulate neural development (Yapijakis, 2020) and different steps of NC formation (Antonaci and Wheeler, 2022; Weiner, 2018).

Neural and NC progenitors are induced in the dorsal ectoderm of vertebrate embryos concomitantly with the positioning of the axial and paraxial mesoderm under the ectoderm. The dorsal ectoderm is patterned into three main territories, the neural plate (NP) in the midline, the adjacent neural border (NB) and the non-neural ectoderm (NNE) laterally (Pla and Monsoro-Burq, 2018). The NP will form the central nervous system, while the NNE participates in skin external layers. Between these two territories, the NB domain includes progenitors for the dorsal-most NP cells, the hatching gland progenitors in frogs and the NC cells and cranial placodes (Betancur et al., 2010; Godden et al., 2021; Pla and Monsoro-Burq, 2018; Steventon and Mayor, 2012). Fate decisions in the dorsal ectoderm are controlled by finely tuned spatial-temporal modulations of BMP, WNT and FGF signalling (Alkobtawi et al., 2021). In particular, the levels of BMP activity, forming a dorsal (low) to lateral (high) gradient, need to be strictly balanced for NB formation. Higher BMP promotes NNE fate while low BMP favours NP, but in both situations the NB derivatives may fail to form (De Robertis and Kuroda, 2004). Low BMP signalling is achieved in the dorsal midline by production of BMP antagonists such as Chordin (Chd1/2 in frog) and Noggin (Plouhinec et al., 2013). Above the paraxial and intermediate mesoderm, slightly higher BMP signals synergize with a complex temporal sequence of WNT and FGF signalling to promote NB formation (Alkobtawi et al., 2021). These signalling modulations result in the activation of key transcription factors in the ectoderm. Specifically, the relative activity of *Pax3/7* and *Zic1* controls fate choices in the dorsal ectoderm. In the NB in particular, finely tuned cooperation between Pax3 and Zic1 promotes NC induction (*snail2, sox10*) (Milet et al., 2013; Monsoro-Burq et al., 2005), higher Pax3 expression triggers hatching gland patterning (*Xhe)*, while higher Zic1 favours cranial placode specification (*six1/3, eya1*) ((Hong and Saint-Jeannet, 2017). In parallel, *Zic3* and *Zic1* genes have an initial pro-neural action during gastrulation, favouring NP specification (*sox2/3)* at the midline (Delaune et al., 2005).

From NB progenitors, pre-migratory NC cells form from gastrulation to neurulation and end up at or close to the developing neural folds. They exit from the dorsal epithelium by a stereotypical EMT, and migrate towards multiple locations of the embryonic body, where they differentiate into many differentiated cell types and tissues. The NC forms neurons and glia of the peripheral sensory, autonomous and enteric nervous system, the craniofacial skeleton and mesenchyme, the adrenal medulla chromaffin cells, and pigment cells (Aoto et al., 2015; Cheung and Briscoe, 2003). These discreet fates depend on the origin of the cells along the body axis, on the diversity of programs elicited prior to emigration, on signals encountered along the migration paths and on niches and environmental signals at the destination (Sauka-Spengler and Bronner-Fraser, 2008). Any errors in these processes can result in neurocristopathies, a large group of congenital disorders where aberrant NC migration, specification, or differentiation leads to multiple defects in NC derivatives (Gouignard et al., 2016; Ward et al., 2018). Since NC development from its induction to its differentiation is orchestrated by evolutionarily conserved gene regulatory networks (NC-GRN), a better understanding of the molecular mechanisms leading to neurocristopathies can be reached using cellular models and animal models (Medina-Cuadra and Monsoro-Burq, 2021; Wulf et al., 2021). Some neurocristopathies correlate with dysregulation of miRNA expression, suggesting that miRNAs may be therapeutic targets for these disorders (Bachetti et al., 2021; Evsen et al., 2020; Pilon, 2021; Schoen et al., 2017). We previously characterised the expression of miRNAs in early *Xenopus* development (Ward et al., 2018) and examined miRNAs that potentially play a role in NC development (Godden et al., 2021; Godden et al., 2023; Ward et al., 2018). Here, we find that miR-196a is required for NC development *Xenopus laevis* embryos, and that its depletion in cranial NC strongly impairs craniofacial skeleton formation.

## RESULTS AND DISCUSSION

### miRNA 196a is essential for neural crest-derived craniofacial development

We previously characterised miR-196a expression in *Xenopus* embryos during late gastrula and neurula stages (Godden et al., 2021; Ward et al., 2018). To assess miR-196a function in NC development two antisense morpholino (MO) oligonucleotides were designed, one complementary to the mature *Xenopus laevis* miRNA (miR-196a-MO), the other as a mismatch control containing 5-mismatched residues (miR-196a-MM, Table 1). MO binding to the miRNA prevents it from interacting with its target mRNAs (Flynt et al., 2017). The miR-196a-MO efficiently decreased miR-196a expression in a dose-dependent fashion as shown by qPCR while the miR-196a-MM had no effect, compared to un-injected control samples (Fig. S1A). MO miR-196a-MO is complementary to both miR-196a and -b isoforms, however we showed effective rescue of miR-196a KD with addition of miR-196a mimic, but we do not rescue loss of miR-196b, therefore the rescue experiment is isoform specific to miR-196a (Fig. S1D-F). Injection of miR-196a-MO leads to craniofacial phenotypes, although mild, that can be seen externally on stage 28 tadpoles, with asymmetric branchial arches (Fig. 1B). Moreover, miR-196 downregulation caused by miR-196a-MO was rescued by co-injection of a *Xenopus tropicalis (xtr)* miR-196a mimic, a synthetic RNA oligonucleotide mimicking xtr-miR-196a (Fig. S1D-G). The miR-196a mimic alone did not have a developmental phenotype (Fig. S1C). Together these data demonstrate the efficiency and the specificity of our *in vivo* depletion strategy.

**Figure 1:**
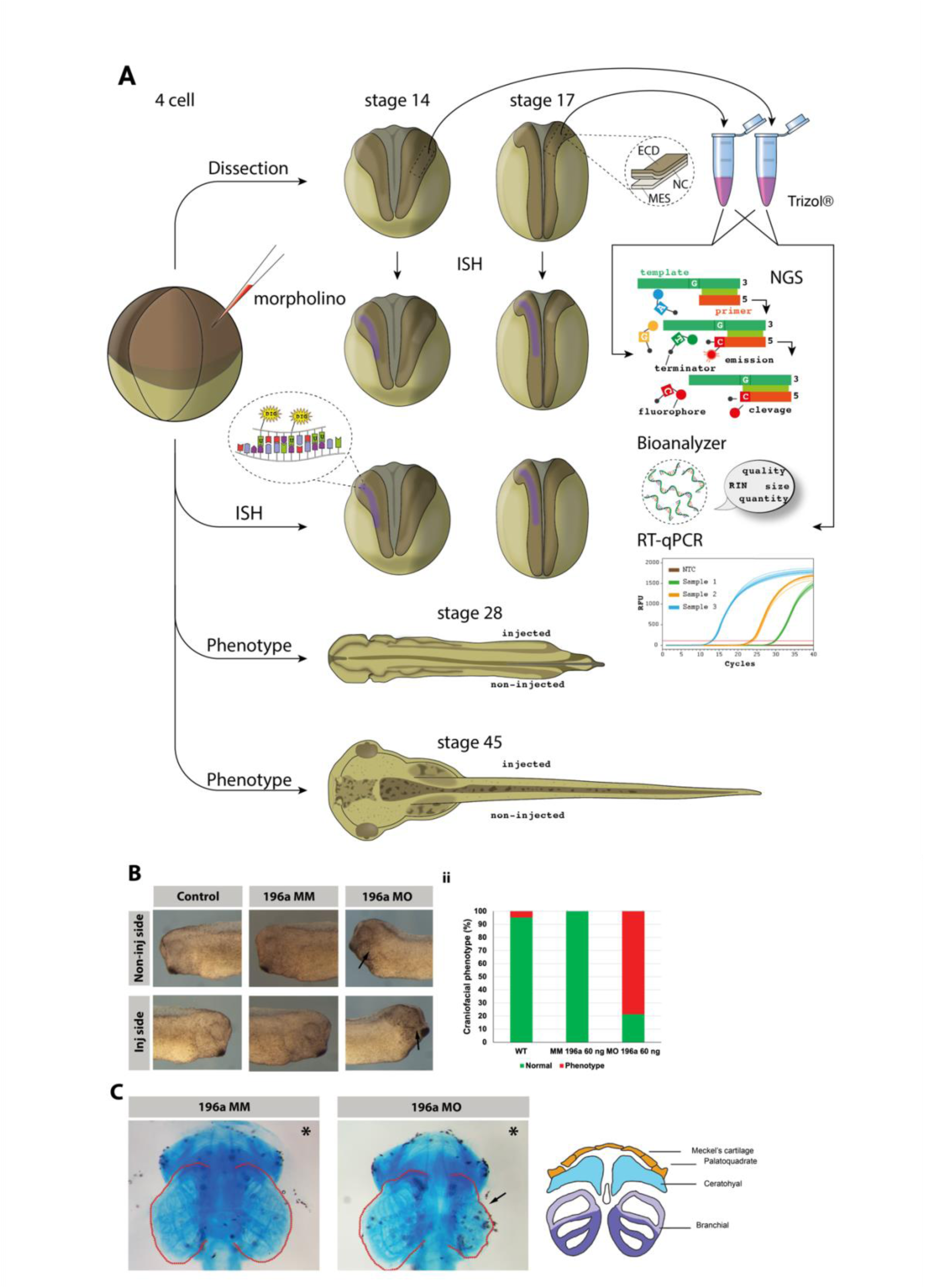
Depletion of miR-196a impairs neural crest-derived craniofacial development. (A) Experimental paradigm for RNA sequencing of single NB or NC explants. Embryos were injected into one dorsal blastomere at the 4-cell stage of development. All embryos were co-injected with 5 ng GFP capped RNA. (B) Craniofacial phenotypes seen in tadpoles are indicated by arrows (stage 28). The graph shows blind count data for wild type, mismatch miR-196a MO and miR-196a MO. Test used for statistical analysis is *Chi*-squared on independent repeats. WT vs miR-196a MO *p* = 0.006, miR-196a MM vs MO *p* = 0 .033. (C) Alcian blue cartilage preparations show clear branchial arch and cartilage phenotypes on the injected side following miRNA-196a KD (stage 45).

**Table 1:**
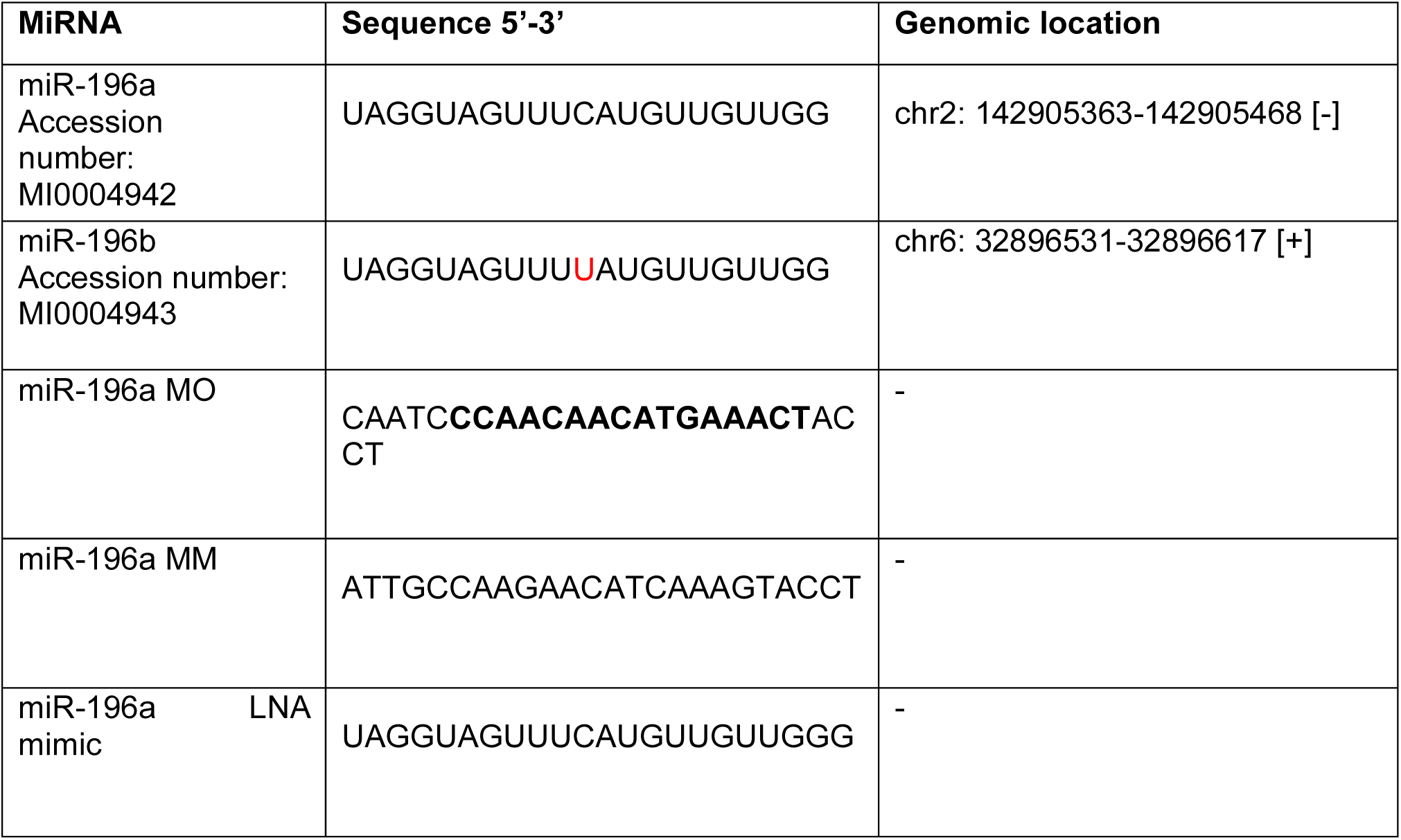
Comparison of miR-196a vs miR-196b sequences in relation to designed MO. Sequences obtained from miRbase (http://www.mirbase.org/). Mature miRNA sequence is highlighted in bold.

We next assessed the developmental phenotypes resulting from unilateral depletion of miR-196a, the various approaches used are illustrated in Fig. 1A. At stage 28, control tadpoles present a well-developed head and facial bulges where the NC populates the branchial arches (Fig. 1B arrows). On the injected side, both control and miR-196a-MM-injected tadpoles formed craniofacial structures normally. In contrast, the tadpoles injected with miR-196a-MO lacked the branchial mesenchyme structures (Fig. 1B) as observed when NC cells are ablated (Milet et al., 2013). The other parts of the larvae were overall well developed, albeit a small size reduction (Fig. 1, Fig. S1B). This phenotype remained until the end of embryonic development, indicating that it was not due to a delay in craniofacial morphogenesis. Alcian blue-stained cartilages of stage 42 morphant tadpoles showed a phenotype on the injected side, with reduced and poorly differentiated cartilaginous branchial arches structures (Fig. 1C, arrow), and missing Meckel’s and palatoquadrate cartilages. Midline skeletal structures remained unchanged. In other parts of the body, NC-derived melanocytes and fin mesenchyme seemed normal (Fig. S1C). These results match CRISPR/Cas9 knockdown of miR-196a (Godden et al., 2021) and indicate that miR-196a plays a major role in NC development, especially for its ectomesenchymal cranial derivatives, prompting us to explore further at which stage the NC-GRN may be affected.

### miRNA 196a regulates dorsal ectoderm patterning

To investigate at which developmental stage miR-196a affected the development of NC or its surrounding tissues, whole mount *in situ* hybridisation for the main early markers of each fate was carried out on unilaterally injected morphant embryos (Fig. 2). At NP stage (stages 13-15) or neural fold stage (stages 16-18), depletion of miR-196a led to a loss of expression for the NC markers *snai2, sox10* and the NB marker *pax3* while miR-196a-MM did not affect gene expression compared to the control un-injected side (Fig. 2A, Fig. S2). Interestingly, *pax3* expression in the superficial ectoderm of the hatching gland seemed unaffected (Fig. 2A, white arrow) while expression in the underlying dorsal neural tube and NC was reduced. This indicated that miR-196a, expressed from late gastrula/early neurula stages, affected dorsal early neurula ectoderm patterning. We further explored neural and NB formation. We found that NP (*sox2, sox11)* and NP/NB markers (*zic1/3)* were expanded, while the lateral NB marker *msx2* was absent from anterior NB (Fig. 2B, Fig. S2). *Pax6* was slightly expanded in the eye progenitor domain and reduced in the NP territory, while notch-responsive *Hairy1* gene was decreased in both domains (Fig. 2B). miR-196a is located within an intron of the hoxC9 gene (Godden et al., 2021). Anterior-posterior patterning was therefore probed using *engrailed2 (en2)* which marks the mid-hindbrain boundary: either a posterior shift or lack of expression was observed in morphant neural tissue (Fig. 2B, Fig. S2C), suggesting an anteriorisation of the morphant neural domain at neural fold stage. This is consistent with the observed expansion of *pax6-*positive eye domain and recess of *pax6* spinal cord domain (Fig. 2B). Importantly, the specificity of these phenotypes was confirmed by rescuing the MO phenotype using injection of miR-196 mimic on *snai2, sox10 (NC)* and *pax3* (NB) (Fig. S2A-B). For all markers no phenotype was seen following injection of embryos with either control miRNA mimic (cel-miR-39-3p), miR-196a mimic, or mismatch 196a-MM alone (Fig. S2A-B). Collectively, these results indicate perturbation of dorsal ectoderm patterning upon depletion of miR196a. Imbalanced expression of essential NB specifiers is known to result in failure of NC formation in favour of adjacent neural or non-neural tissue development (de Croze et al., 2011). Here we observe a failure of activation for the key transcription factors of the NB-NC-GRN *pax3, msx2, snai2,* and *sox10*. Neural progenitors are expanded, especially anteriorly (marked by *sox2, sox11, pax6, zic1/3),* however, we observed that neuronal differentiation was impaired at the end of neurulation as differentiation markers *tubb2b (*formerly *N-tub) and elrC* were decreased on the morphant side (Fig. 2B).

**Figure 2:**
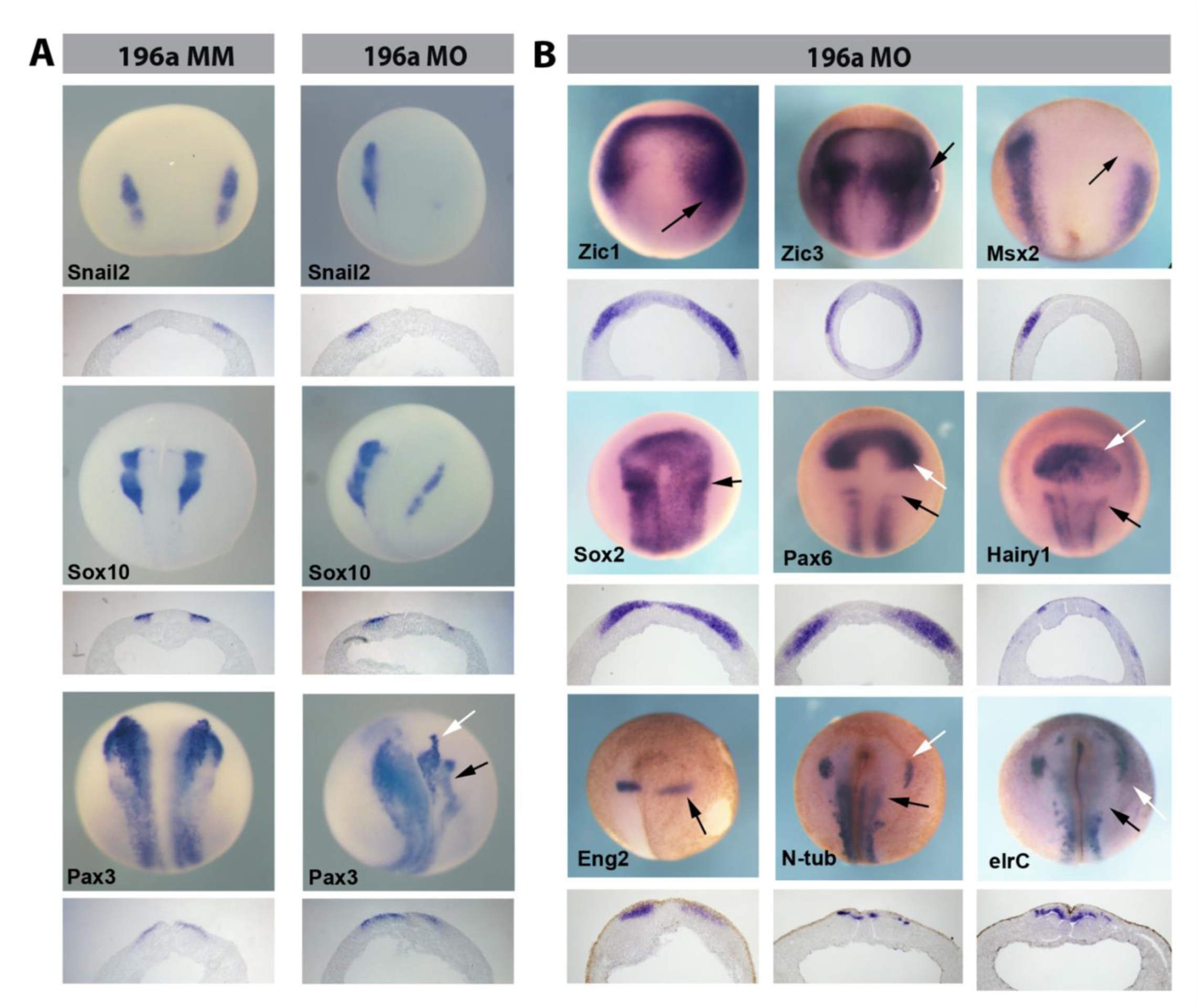
Depletion of miR-196a expands neural plate progenitors at the expense of neural crest specification and neuronal differentiation. Embryos were injected into one dorsal blastomere at the 4-cell stage of development with 300 pg of GFP mRNA or of lacZ developed with Red-gal. In all panels the injected side is to the right. (A) miR-196a MO KD and WISH for *snail2* (stage 14), *sox10* (stage 14) and *pax3* (stage 15), with control groups: miR-196a MM and miR-196a MO. (B) Assessment of neural, NP and placodal development following MO mediated miRNA-KD. Embryos were injected into one dorsal blastomere at the 4-cell stage of development with MO and 300 pg of lacZ developed with Red-gal. (B) Whole mount *in situ* hybridisation of *zic1*, (stage 14), *zic2*, (stage 14), *msx2*, (stage 14), *sox2*, (stage 14), *pax6*, (stage 15) *hairy1*, (stage 18), *eng2*, (stage 15) *n-tub* (stage 18) and *elrc1,* (stage 18), following MO mediated miRNA KD. Wholemounts are shown as well as transverse sections.

### Transcriptomes of mir-196a morphant NC progenitors indicate loss of NC and placodes, expansion of immature neural programs and aberrant BMP/Notch signalling

Using RNA sequencing, we compared micro-dissected NB at NP stage (stage 14) with premigratory NC at neural fold stage (stage 17), after 196-MO or 196-MM injections (Fig. S3). The ectoderm layers were taken, while the underlying mesoderm was avoided, as in (Plouhinec et al., 2017). Accuracy of the dissection was tested by probing the donor embryos with *pax3* for NB and *snai2* for NC, after dissection (Fig. S3A and B). PCA analysis of the triplicates indicates grouping of the samples according to their injection type (Figs. S3C and D). In addition, correlation analysis confirmed that the transcriptomes of control samples were grouped according to developmental stage and together, while the morphant transcriptomes were grouped separately (Fig. 3A). Interestingly, stage 17 morphant samples grouped closer to the control stage 14, raising the possibility that the phenotype could resemble a developmental delay. However, our phenotype analysis (Fig. 1) indicates that any potential delay is not compensated later, as also shown in other experimental settings where NC development is delayed during neurulation (Figueiredo et al., 2017).

**Figure 3:**
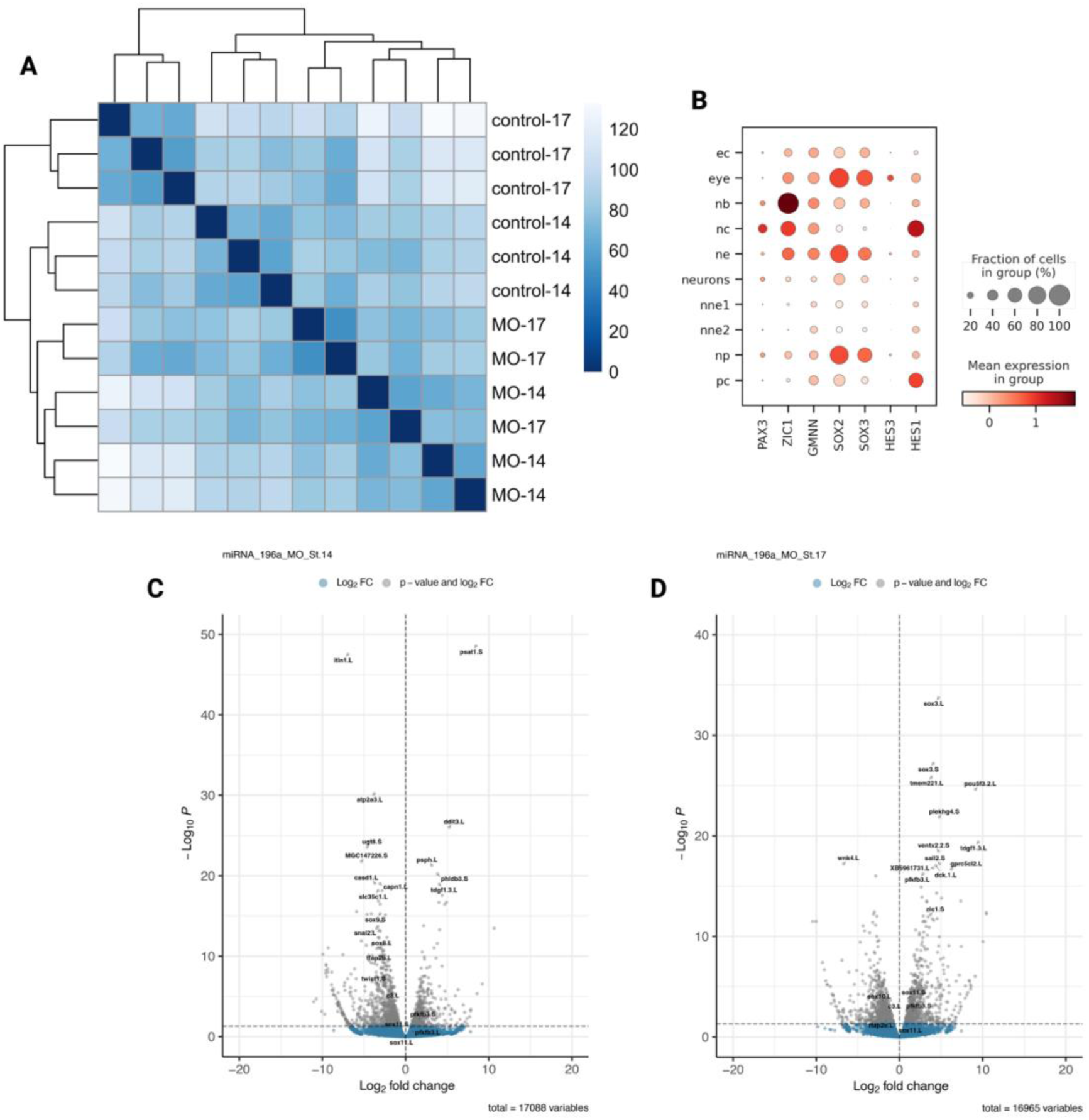
Main features of RNA sequencing of neural border and neural crest after depletion of miR-196a. (A) Heatmap showing clustering of control and miR-196a KD samples. (B) Dotplot of NPB, neural and NC gene expression across tissues. Ec, ectoderm; nb, neural plate border; nc, neural crest; ne, neuroectoderm; nne, non-neural ectoderm; np, neural plate; pc, placode. (C-D) Volcano plots of differentially expressed genes follow miR-196a MO KD at stage14 (C) and stage17 (D).

We used tissue expression signatures from single cell data to define the expression profile of landmark genes (Kotov et al., 2024; Petrova et al., 2024). A dot plot represents the intensity and percent of positive cells expressing a given gene in a cell type at different developmental stages (ectoderm, eye, NB, NC, neural ectoderm, neurons, NNE, NP and placodes) and illustrates the promiscuity of gene expression (Fig. 3B). Among the 2432 genes differentially expressed in NB explants at NP stage 14, the NC markers *snai2, sox8, sox9, tfap2b, c3 and twist1* were depleted (Fig. 3C) while at that stage, NP markers remained little changed (e.g. *sox2, zic1).* In contrast, at the end of neurulation, in NC explants that should be ready to undergo EMT and cell migration, the morphant NC cells exhibited aberrant high expression of early NP markers *sox2/3* and of pluripotency genes *ventx2.2* (nanog ortholog) and its target *pou5f3 (oct4* ortholog) (Fig. 3D). Those genes are essential for earlier steps of NC progenitor formation but should be downregulated at that stage (Scerbo and Monsoro-Burq, 2020).

We further explored ectoderm patterning processes using significantly differentially expressed genes from RNA-seq data and comparing early and late neurulation stages (Fig. 4B, Fig. S4B). Expression of NB specifiers was unbalanced with increased expression of *zic1* or *hes4 (hairy2a)* at both stages, while early *msx1/2* expression and *tfap2b, c and e* levels at both stages were decreased. NC specifiers *snai1/2, sox8, myc, foxd3, ets1*, and the EMT/migration marker matrix metalloprotease, *mmp28*, were not activated in morphant NC cells at either stage. The same profile was observed for cranial placode markers *six1/4, eya1, pax8* and specified NNE markers *gata2/3*, and ectoderm differentiation marker *keratin.* This indicates that the three NB fates, NC, NNE and placodes, which should be clearly identified from NB explants at stage 14 (Kotov et al., 2024) are not observed in miR196a morphant NB. In contrast, early neural development (*sox2/3, zic2/3/4)* and pluripotency gene Oct4 orthologs (*pou5f3.1/2/3)* were highly expressed in NB ectoderm as well as in pre-migratory NC cells.

**Figure 4:**
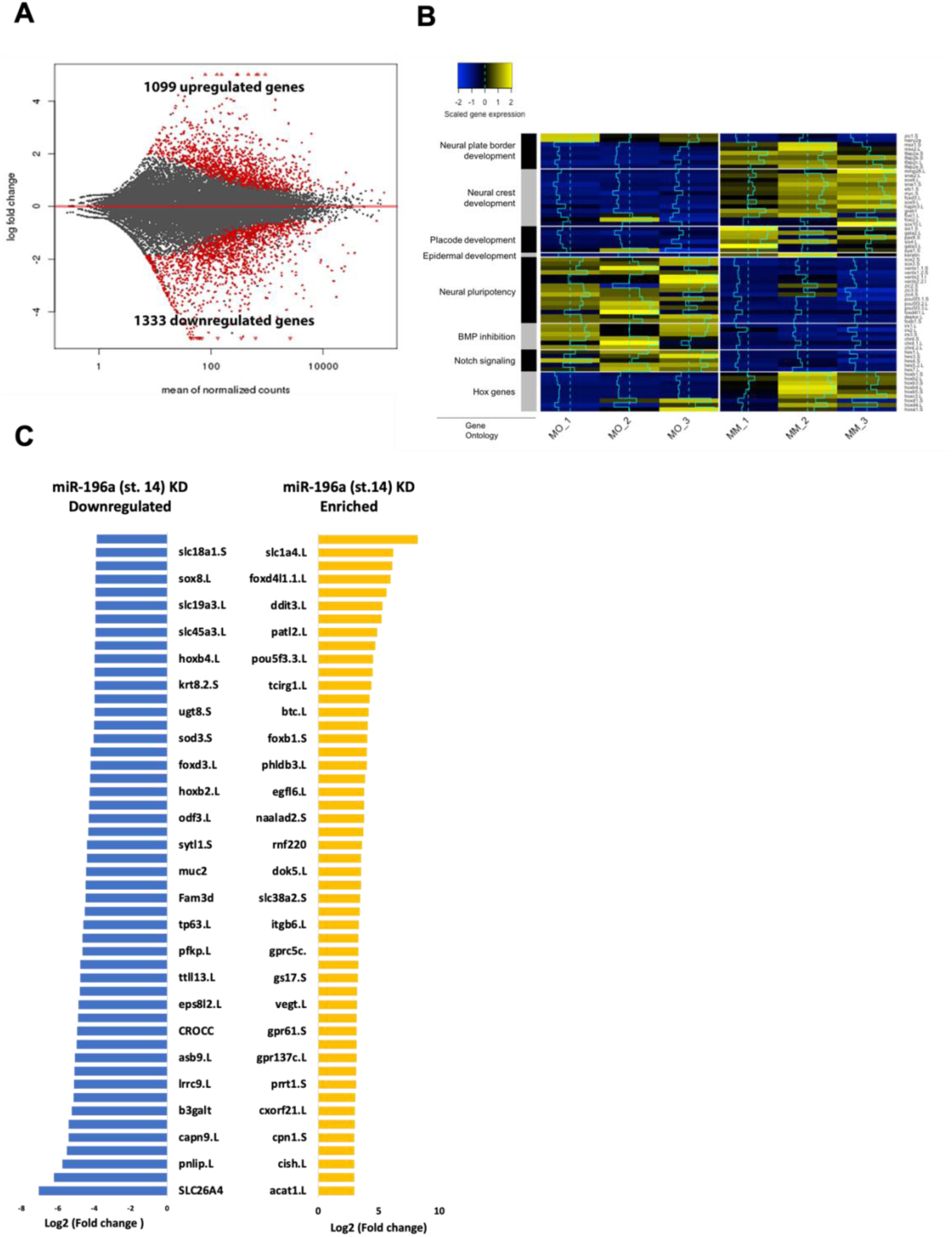
Ectoderm patterning, BMP and Notch signalling are profoundly altered by miR196a depletion. (A) MA plot of significantly differentially expressed genes in RNA-seq data following miR-196a KD (stage 14). (B) Heatmap for selected differentially expressed genes (miR-196a MO vs miR-196a MM). Colour depicts gene expression where yellow represents overexpression and blue under expression. The blue line running throughout the boxes is a histogram representing the level of either upregulation or downregulation against the dotted midline (no change). (C) Top 50 enriched and depleted significantly differentially expressed genes following miR-196a MO KD at stage14.

**Figure 5:**
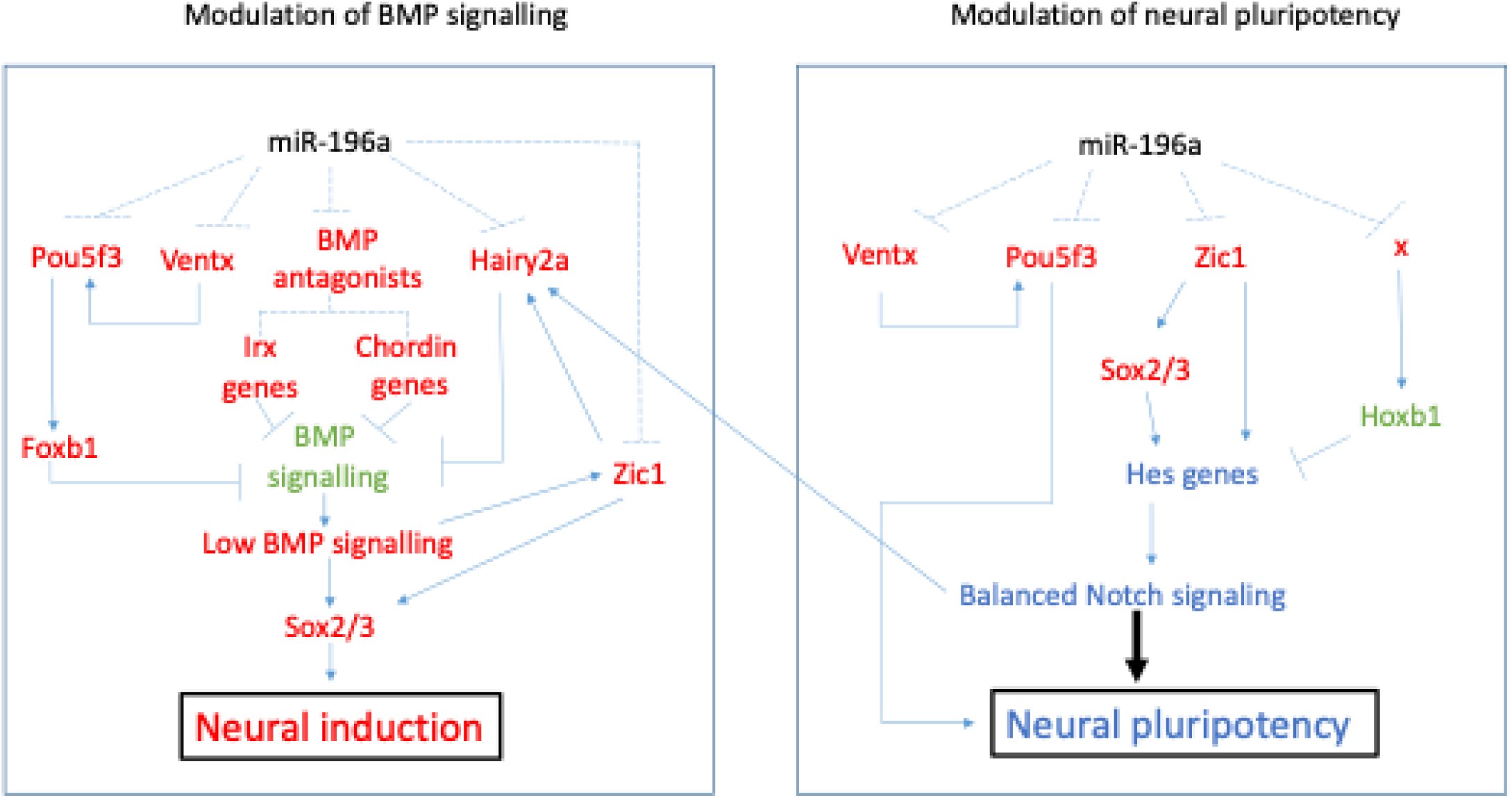
Model of the potential molecular mechanisms miR-196a uses to prevent neural induction and control neural pluripotency during Xenopus neuroectoderm patterning. Red represents repression, green represents expression and blue represents balanced. Solid and dashed lines are the verified and predicted regulatory relationships, respectively.

As BMP signalling balance is critical for dorsal ectoderm patterning, we were puzzled to observe chordin (*chrd)* expression in those explants, which do not contain mesoderm as confirmed prior to sequencing by checking for mesoderm genes. Chordin, however, is expressed in the floor plate ectoderm, which shares common embryonic origins with the notochord (Le Douarin et al., 2004). This expression thus must be controlled negatively in other ectoderm area during normal development. Our observation indicates that upon KD of miR196a, BMP antagonist Chordin expression is potentially repressed in the NB progenitors, creating a lower BMP signalling level than the one required for NB patterning (Alkobtawi et al., 2021). By itself, this result could explain the set of ectoderm patterning defects observed, with enlarged immature neural tissue and loss of NB derivatives.

Furthermore, pluripotency factors (*ventx2.2, pou5f31/2/3)*, neural stemness genes such as *Sall3* (Kuroda et al., 2019) and Notch signalling effectors (*hes3/4/5/1)* regulate the balance between progenitor state and differentiation (Nichane et al., 2008a; Nichane et al., 2008b). Depletion of mir-196a led to maintained expression of these genes in the NB and NC tissues during neurulation, in agreement with the expression patterns observed in toto (Fig. 2). Last, anterior-posterior patterning of the NC, as well as of the adjacent neural tube, is largely regulated by *hox* genes posterior to the mid-hindbrain boundary. In the morphant NB explants, which include both anterior and posterior ectoderm, increased *hox* gene expression is observed compared to control tissue. However, in the stage 17 pre-migratory NC explants, which normally include both anterior *hox*-negative cells and *hoxa-b-c-d1/2/3-*positive rhombencephalic cells, miR-196a depletion eliminated hox expression in the morphant tissue (Fig. 4, Fig. S4). It remains unclear, at this tissue resolution if this is due to loss of NC cells, or to their transformation into a mis patterned immature neural progenitor state.

Among the genes mis-regulated in morphant tissue, we hypothesized that some might be direct miRNA targets, which should be upregulated upon miRNA depletion, while others could be affected indirectly. We collated genes which are predicted to be miR-196a targets, using miRanda with default settings, (John et al., 2004). These include NC genes *sox10*, *hand1* and Hox genes *hoxc8, hoxd8, hoxb3, hoxb6* and neural and ectoderm gene *pax2*, and placodal gene *eya1* (Table 2). To directly validate this potential regulation *in vivo* during ectoderm patterning, we then tested *in toto* expression of *sox10* in unilateral miR-196a KD and rescue injections (Fig. S2A and B). Surprisingly the expression of *sox10* was reduced after MO KD and rescued upon co-injection with miRNA mimic. However, we did observe a striking expansion for the neural and placode marker *sox11*, encompassing the entire dorsal ectoderm and consistent with the morphant patterns described above (Fig. S2E). *Sox11* is not a direct target of miR-196a in *Xenopus*, however it was found to be a target in humans when analysis TargetScan database. In future annotations of *Xenopus* genomes, it would be worth revisiting this as we found *sox11* to be enriched following miR-196a KD in our RNA-seq data. This analysis confirms that some direct targets for miR-196a are tightly controlled during vertebrate embryo ectoderm, neural and NB patterning.

**Table 2:**
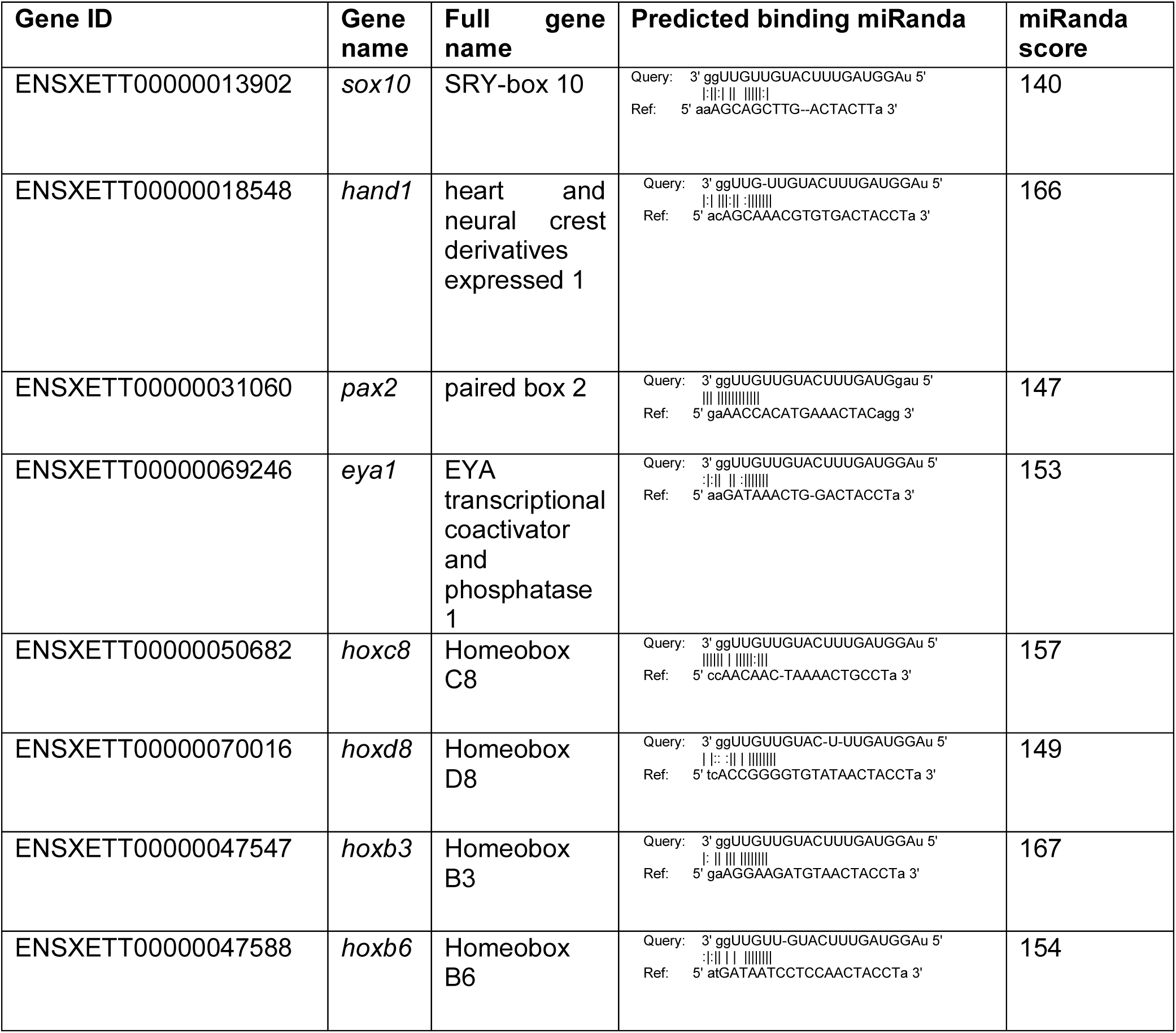
Results of miRanda analysis for miR-196a. Contents cover Gene ID, name and predicted binding of miR-196a (query) and 3’ UTR of gene (Ref), with miRanda score in last column, higher score indicates stronger match.

Overexpression of miR-196a has been shown to cause eye defects in *Xenopus* embryos (Qiu et al., 2009). Work aiming to investigate miR-196a further identified anterior neural development phenotypes (Gessert et al., 2010). Therefore, it was expected that following misexpression of miR-196a with MO or miRNA mimic that there would be phenotypes generated on eye markers and neural markers. As expected, the neural marker Pax6, responsible in eye development (Grocott et al., 2020), is enriched after miR-196a KD. Interestingly the skeletal preparations for miR-196a KD tadpoles showed craniofacial defects (Fig. 1), indicative of abnormal NC specification, this has also been seen in previous work by Gessert and colleagues (Gessert et al., 2010) and our own miR-196a CRISPR knockouts (Godden et al., 2021)

### Conclusion

Collectively, our findings highlight the essential role of miR-196a in patterning to dorsal ectoderm of the vertebrate embryo model *Xenopus laevis*. At the NP stage, when the different ectodermal fates are established, miR-196a activity controls the expression of more than 2,400 genes. These include direct and secondarily regulated genes. Depletion of miR196 was carefully controlled using mimic RNA oligonucleotides *in vivo*, an experimental strategy which was not described before, and allows us to exclude compensatory action of the paralog miR-196b. The main function of miR-196a seems to restrict neural fate in the dorsal ectoderm. When mir196a is depleted, this results in failure to form other ectoderm derivatives such as NC and placodes and NNE. Some tissues, such as the hatching gland ectoderm forming at later neurula stage, independently of the initial dorsal ectoderm patterning, seemed unaffected or even slightly increased in the morphant context. These observations are consistent with *sox2* expansion in miR-196a KD embryos (Fig. 2) and could contribute towards the reduction in NC marker expression seen in Fig. 2; similar to previous observations in avian models suggesting that *sox2* misexpression can inhibit NC formation (Hu et al., 2014; Wakamatsu et al., 2004).

Defective NC formation is not compensated for at later stages of development, while it seems that other early defects in the non-NC tissues are transient since tadpole morphology is globally normal. *Sox10* depletion is known to lead to a rise in NC cell apoptosis and reduction in cell proliferation (Honore et al., 2003). *Snail2* is required for the induction of NC; and is anti-apoptotic (Klymkowsky et al., 2010; Shi et al., 2011). This suggests that miR-196a KD, which reduced expression of *sox10* and *snail2* (Fig. 2), is preventing proliferation and maintenance of a pool of NC cells leading to the craniofacial deformation seen later in development (Fig. 1C). The second major function of miR-196a is to repress expression of pluripotency and neural stemness genes and allow the onset of neuronal differentiation in the central nervous system. We propose a working model (Fig 6) to place miR-196a in the current neural/neural border/non-neural ectoderm gene regulatory network, and a rich resource of candidate genes for further validation of their functions in the initial steps of ectoderm regionalisation in vertebrates.

## MATERIALS AND METHODS

### Xenopus husbandry

All experiments were carried out in accordance with relevant laws and institutional guidelines at the University of East Anglia, with full ethical review and approval, compliant to UK Home Office regulations. To obtain *Xenopus laevis* embryos, females were primed with 100 units of PMSG and induced with 500 units of human chorionic gonadotrophin. Eggs were collected manually and fertilised *in vitro*. Embryos were de-jellied in 2% L-cysteine, incubated at 18°C and microinjected in 3% Ficoll into 1-cell at the 2/4-cell stage. Embryos were left to develop at 23°C. Embryo staging is according to Nieuwkoop and Faber normal table of *Xenopus* development. GFP/LacZ capped mRNA for injections was prepared using the SP6 mMESSAGE mMACHINE kit, 50 pg were injected per embryo.

### Embryo injection

*Xenopus. laevis* embryos were used for all *WISH* (whole mount *in situ* hybridisation) experiments in this project. Embryos were injected using a 10 nL calibrated needle. MO dose was optimized to 60 ng for miRNAs; MO and GFP or lacZ capped mRNA were injected at 4 cell stage of embryo development into the right dorsal blastomere.

*Xenopus* laevis embryos were injected at the 4-cell stage of development into one dorsal blastomere with 300 pg GFP or lacZ plus MO, miRNA mimic or combination. Synthetic LNA miRNA mimic, complementary to the mature miRNA sequence (SFig. 3B). For all markers use of synthetic miRNA mimic were used to rescue phenotypes generated by miRNA MO KD. Use of miRNA mimic and control mimic cel-miR-39-3p alone saw no impact on embryo development. Rescue of the miRNAs is specific with use of relevant synthetic miRNA (Supp Fig 3). MiRNA expression levels were significantly increased and decreased following use of miRNA mimic, and MO. When used together miRNA mimic rescues decreased expression levels by miRNA MO (Fig. S3).

Embryos were injected using a 10 nL calibrated needle. MO dose was optimized to 60 ng for miRNAs; MO and GFP or lacZ capped mRNA were injected at 4 cell stage of embryo development into the right dorsal blastomere.

MiRCURY LNA miRNA mimics were used to replace miRNA in MO miRNA KD rescue. For miR-196a: (Qiagen, 339173 YM00470616-ADA, MIMAT0000226); hsa-miR-196a-5p compatible with xtr-miR-196a sequence: 5’ UAGGUAGUUUCAUGUUGUUGGG. A negative control miRNA mimic recommended by Qiagen was used (Qiagen, 331973 YM00479902-ADA); Negative control (cel-miR-39-3p), sequence 5’UCACCGGGUGUAAAUCAGCUUG.

Chi-squared test for association was used to test phenotype yes or no categories for MO injected embryos to see if there was a relationship between two categorical values. Excel was used to collate and tabulate data. IBM SPSS v25 to carry out Chi-squared test. When describing statistical significance; p<0.05 = *, p<0.01 = **, p<0.001 = ***, p=<0.0001= ****.

### RT-qPCR

Embryos were frozen on dry ice before RNA extraction. Total RNA was extracted from five stage14 Xenopus tropicalis embryos. Embryos were homogenised with a micro pestle and RNA was extracted according to manufacturer’s guidance, Quick-RNA Mini prep plus kit (Zymo, Cat no. R1058). Samples were eluted in 25 µL of nuclease free water; RNA concentration and purity quantified on a Nanodrop 1000 and 1 µL was checked on a 2% agarose gel.

To produce cDNA for q-RT-PCR, miRCURY LNA RT kit was used (Qiagen, Cat No./ID: 339340). 50 ng of RNA was used and kit according to manufacturer’s instructions. cDNA was produced on a thermocycler with the following programme: 42°C for 60 min and 95°C for 5 min. cDNA was diluted 1:40 for q-RT-PCR. cDNA was stored at −20°C. qRT-PCR reactions were set up in 10 µL volume containing 4 µL cDNA, 1 µL primer (in accordance with manufacturer dilute for Qiagen LNA miRNA primer), and 5 µL SybrGreen (Applied Biosystems 4309155). Primers for q-RT-PCR sequences were the following: xtr-miR-196a 5’ – UAGGUAGUUUCAUGUUGUUGG – 3’ (Qiagen, YP02103491), xtr-miR-196b 5’-UAGGUAGUUUUAUGUUGUUGG – 3’ (Qiagen, YP02104328), and U6 snRNA 5’-CTCGCTTCGGCAGCACA – 3’.

### In situ hybridization

WISH with LNA probes was carried out according to (Ahmed et al., 2015; Antonaci et al., 2023; Sweetman et al., 2006) Other WISH and probe synthesis were carried out according to (Monsoro-Burq, 2007; Sive et al., 2007).

### RNA sequencing

For RNA-sequencing embryos were injected into one blastomere at 4-cell stage with one of two MO’s (miR-196a or miR-196a mismatch (MM)) into one dorsal blastomere to target neural and NC tissue in one side of the embryo. Embryos were left to develop until stage 14 or 17. One group underwent WISH to check NC that genes were knocked down (data not shown) and the other group underwent NC dissections and RNA extraction. Three replicates were collected for each condition (MO, MM, and non-injected control). RNA samples underwent quality control using Bioanalyzer (Agilent) and RT-qPCR was used to further validate the KD of NC-specific genes in MO-injected samples (Godden et al., 2021). Triplicates were then processed to Illumina HiSeq 2500RNA sequencing after unstranded library preparation (50bp paired-end sequencing on the HiSeq High Output run mode PE100 for sequencing.

### Data analysis

RNA-sequencing reads were mapped to the *Xenopus laevis* v10.1 genome assembly using STAR (v.2.7.3a), (Dobin et al., 2013). Differential expression analysis was carried out using DESeq2 (v.1.32.0), (Love et al., 2014) in R (v.4.1.1). Genes with an adjusted *p*-value below 0.05-0.15 were considered significant and were reported by the workflow. The gene model used in the DE bioinformatic analysis was *Xenopus laevis* (NCBI v10.1). For GO enrichment analysis of a DE genes we used ClusterProfiler (v.4.0.5). MiRanda was used to analyse miRNA-196a 3’ UTR gene targets with default settings (John et al., 2004). Mature miRNA sequences were accessed from miRbase (Griffiths-Jones et al., 2006), and annotation and reference fasta file accessed from Ensembl v106 *Xenopus tropicalis* v9.1 genome as miR-196a, b are not annotated in the *Xenopus laevis* genome, but is highly conserved. RNA-sequencing data is available at GEO under accession number: GSE289705 (under embargo until acceptance for publication).

### Alcian blue craniofacial cartilage staining

Stage 45 Tadpoles were dehydrated in ethanol and were then left in Alcian blue (20 mg) for 3 nights. After this, embryos were washed 3 times for 15 min in 95% ethanol (Sigma, UK) then rehydrated in 2% KOH using 10-min washes of 75% ethanol in 2% KOH, 50% ethanol in 2% KOH, 25% ethanol in 2% KOH then 3 x 2% KOH washes. Embryos were then stored in glycerol with 1-hour washes of 20% glycerol in 2%, 40% glycerol in 2% KOH, 60% glycerol in 2% KOH and finally stored in 80% glycerol in 2% KOH. Embryos were washed 3 times for 5 mins in PBS before 20 min incubation in 10 mL pre-incubation solution (0.5 X SSC (150 mM NaCl, 15 mM sodium citrate, pH 7.2), 0.1% Tween. 20), and then embryos were incubated in 10 mL of depigmentation solution (5% formamide, 0.5 X SSC, 3% H2O2). Embryos were then carefully dissected under a microscope using fine forceps to remove outermost ectoderm and mesenchyme surrounding craniofacial cartilage. Method for this clearing was based on (Affaticati et al., 2018).

Embryos were imaged on agarose dishes with a Zeiss Axiovert Stemi SV 11, Jenoptik ProgRes C5 camera (Germany), ProgRes software version 2.7.6. Fluorescent images were captured using Leica MZ 16 F microscope, Leica DFC300 FX camera, Leica Kubler codix light source, Leica FireCam software version 3.4.1.

## ACKNOWLEDGEMENTS

We would like to thank Prof. Saint-Jeannet and Dr. Walmsley for plasmids, Prof. Münsterberg and Prof. Simon Moxon for helpful discussions and the Wheeler, Münsterberg, and Monsoro-Burq labs for support. We would also like to thank Johannes Wittig for drafting Fig. 1A.

This work was supported by the UKRI Biotechnology and Biological Sciences Research Council Norwich Research Park Biosciences Doctoral Training Partnership [Grant numbers BB/M011216/1 to GW/AG] <u>and BB/J014524/1 to NW/GW).</u> This project also received funding from European Union Horizon 2020 Marie Skłodowska-Curie grant no. 860635, NEUcrest ITN (AK, MA, AHMB, GW); Agence Nationale pour la Recherche ANR-15-CE13-0012-01, ANR-21-CE13-0028; Institut Universitaire de France (AHMB); Fondation pour la Recherche Médicale (AHMB). RNA sequencing used the ICGex NGS platform of Institut Curie (ANR-10-EQPX-03; ANR-10-INBS-09-08; Canceropole Ile-de-France; SiRIC-Curie INCa-DGOS-4654)

## AUTHORS CONTRIBUTIONS

GW and AMB designed the study and directed the project. AG, NW, MS, MA and AMB performed the experiments and acquired the data. AG, NW and AK analysed the bioinformatics data. GW, AMB, NW and AG analysed the data. AG, GW and AMB and wrote the manuscript.

## SUPPLEMENTARY MATERIALS

**Supplementary Figure 1-MO and miRNA mimic validation experiments**. (A) q-RT-PCR validation of MO dose-response to show MO specificity. One-way Anova with post-hoc Tukey tests show statistical significance, with p=<0.05 for “*” as the threshold, p=<0.01 “**” and p=<0.001 for “***”, and ns= not significant. (B-C) miRNA mimic, and MO response, used 300 pg of lacZ or GFP capped RNA as a tracer. No phenotypes were observed in the tadpoles (B - stage 28, C-stage 39). (D) q-RT-PCR validation of miRNA levels following miRNA mimic overexpression, miRNA MO, and then rescue using miRNA MO and miRNA mimic. For miR-196a rescue: miR-196a mimic 15 µM vs miR-196a MO 60 ng *p* = 0.01, miR-196a MO vs miR-196a MO 60 ng + miR-196a mimic 15 µM *p* = 0.007, miR-196a mimic 15 µM vs miR-196a mimic 15 µM + miR-196a MO 60 ng *p* = 0.035. (E-F) Analysis of the specificity of miR-196a MO on the miR-196b isoform. MiR-196a MO has capacity to knockdown both miR-196a and miR-196b but is more specific to miR-196a with a 92% reduction in expression compared to an 81% reduction in expression MM vs MO miR-196a vs miR-196b. Percentages show MM vs MO above MO column, the MO vs rescue above the rescue column. Error bars show mean +/- S.E.M, statistical significance measured by T-test. P values: for miR-196a mismatch control vs MO *p* = 0.003, for MO vs rescue *p* = 0.007. For miR-196b p values: miR-196b on miR-196a MM vs MO *p* = 0.006, for MO vs rescue *p* = 0.08. Carried out with three biological and three technical replicates. snU6 housekeeping gene was used for normalisation. MM= mismatch morpholino and MO= morpholino, rescue= MO 60 ng + miR-196a mimic 15 µM. Error bars depict mean± S.E.M with three biological and technical triplicates. Embryos were injected into both blastomeres at 2-cell stage. (G) MiRNA sequences showing alignment of miRNA with mature miRNA, miRNA mimic used to rescue phenotypes, MO sequence, and see region of the miRNA.

**Supplementary Figure 2 - Functional characterisation of MO mediated miRNA KD of miR-196a with synthetic miRNA mimic rescues and assessment of, neural crest, neural, neural plate and placodal development following MO mediated miRNA-KD** Embryos were injected into one dorsal blastomere at the 4-cell stage of development with 300 pg of lacZ developed with Red-gal. (A) miR-196a MO and rescue for *snail2*, (stage14), *sox10*, (stage 14), *pax3* (stage 14), and *xhe2* (stage 18),with control groups: Control miR-mimic, MM MO, miR-196 mimic and miR-196a MO and control miR-mimic. (B) Count data for phenotypes for each marker gene. *Snail2* was carried out with 3 individual biological experimental repeats, statistical analysis was performed using *Chi*-squared 196a MM vs 196a MO *p* = 4.73 x 10^-14^, 196 MO vs rescue *p* = 0.000013. Overall reduction in NC markers was seen along with deregulation of *pax3* and *xhe2* following miRNA KD. Relevant miRNA mimics successfully and specifically rescued phenotypes. Embryos were injected into one dorsal blastomere at the 4-cell stage of development with MO and 300 pg of lacZ developed with Red-gal. (C) WISH of *en2*, (stage 18) *sox2*, (stage 13), *pax6* (stage 14), and *zic1* (stage 13) following MO mediated miRNA knockdown. (D) Blind score phenotype counts. *En2* and *pax6* loss of expression was observed after miR-196a KD with *sox2* and *zic1* expanded. (E) Sox11 following loss of miR-196a showed an expanded phenotype, *p* = 5.9 x 10 ^-77^. MM= mismatch, MO= Morpholino.

**Supplementary Figure 3 - WISH of dissected embryos and PCA plots of RNA-seq samples.** *WISH* was completed on dissected embryos to check dissection accuracy for *pax3* (stage 14) (A) and *snai2* (B) (stage 14). Principal components analysis of sample clustering following morpholino KD of miR-196a in stage14 (C) and stage17 (D) embryos.

**Supplementary Figure 4 - At the end of neurulation miR-196a depletion maintains NC tissue in an immature and posteriorized (Hox-positive) neural state.** (A) MA plot of significantly differentially expressed genes in RNA-seq data following miR-196a KD (st. 17). (B) Heatmap for selected differentially expressed genes (miR-196aMO vs miR-196a MM). Colour depicts gene expression where yellow represents overexpression and blue under expression. The blue line running throughout the boxes is a histogram representing the level of either upregulation or downregulation against the dotted midline (no change), (stage17). (C) Top 50 enriched and downregulated significantly differentially expressed genes following morpholino miR-196a KD at stage17.

## ABBREVIATIONS

EMT: epithelial-to-mesenchymal transition
HG: hatching gland
MO: morpholino miRNA - microRNA
MM: mismatch morpholino
NB: neural border
NC: neural crest
NNE: non-neural ectoderm
NPB: neural plate border
WISH: whole mount *in situ* hybridisation

