## Supplementary figures and images for "MicroRNA miR-196a controls neural crest patterning by repressing immature neural ectoderm programs in Xenopus embryos"

### Supplemental Figures

#### Supplemental figures

##### Supp fig 1

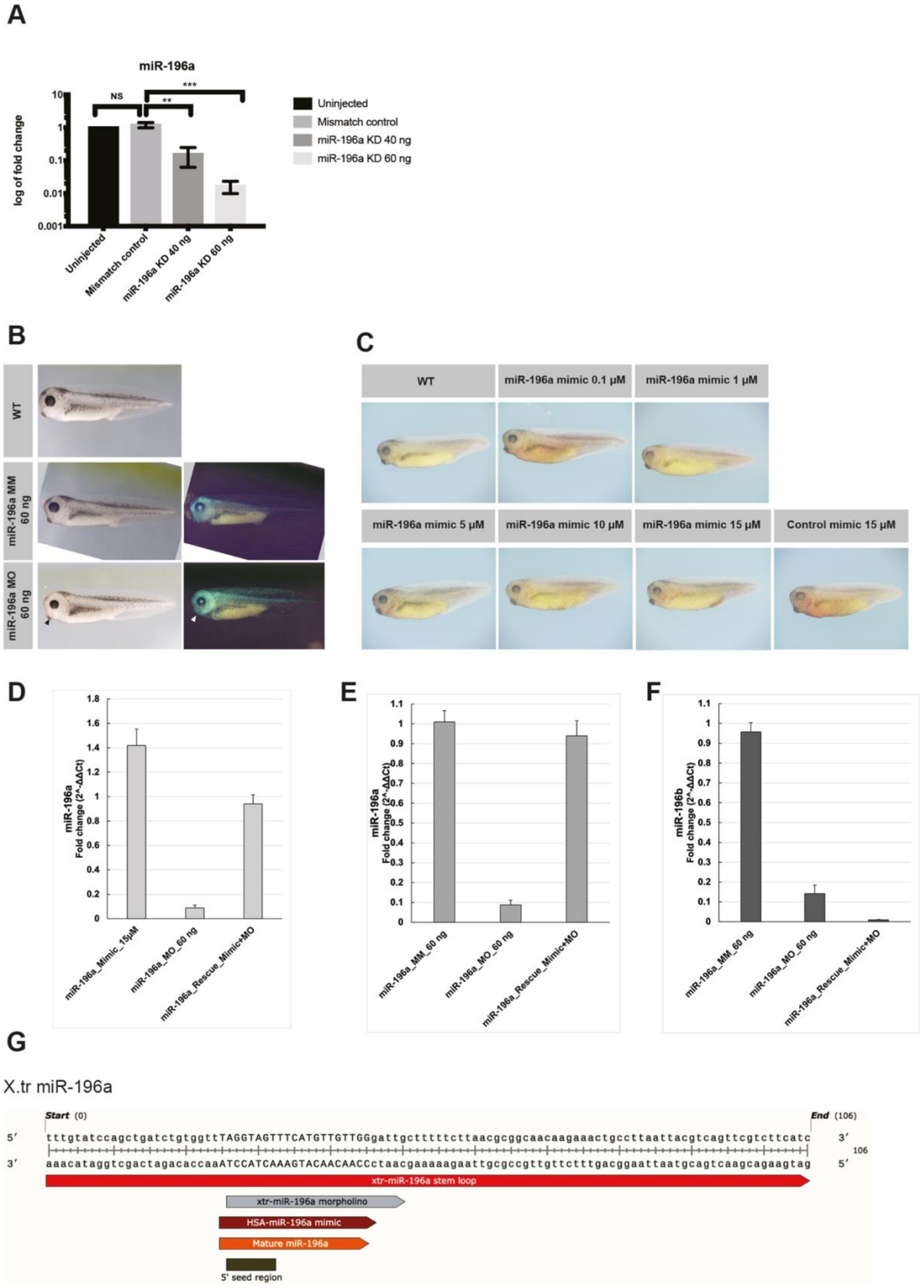

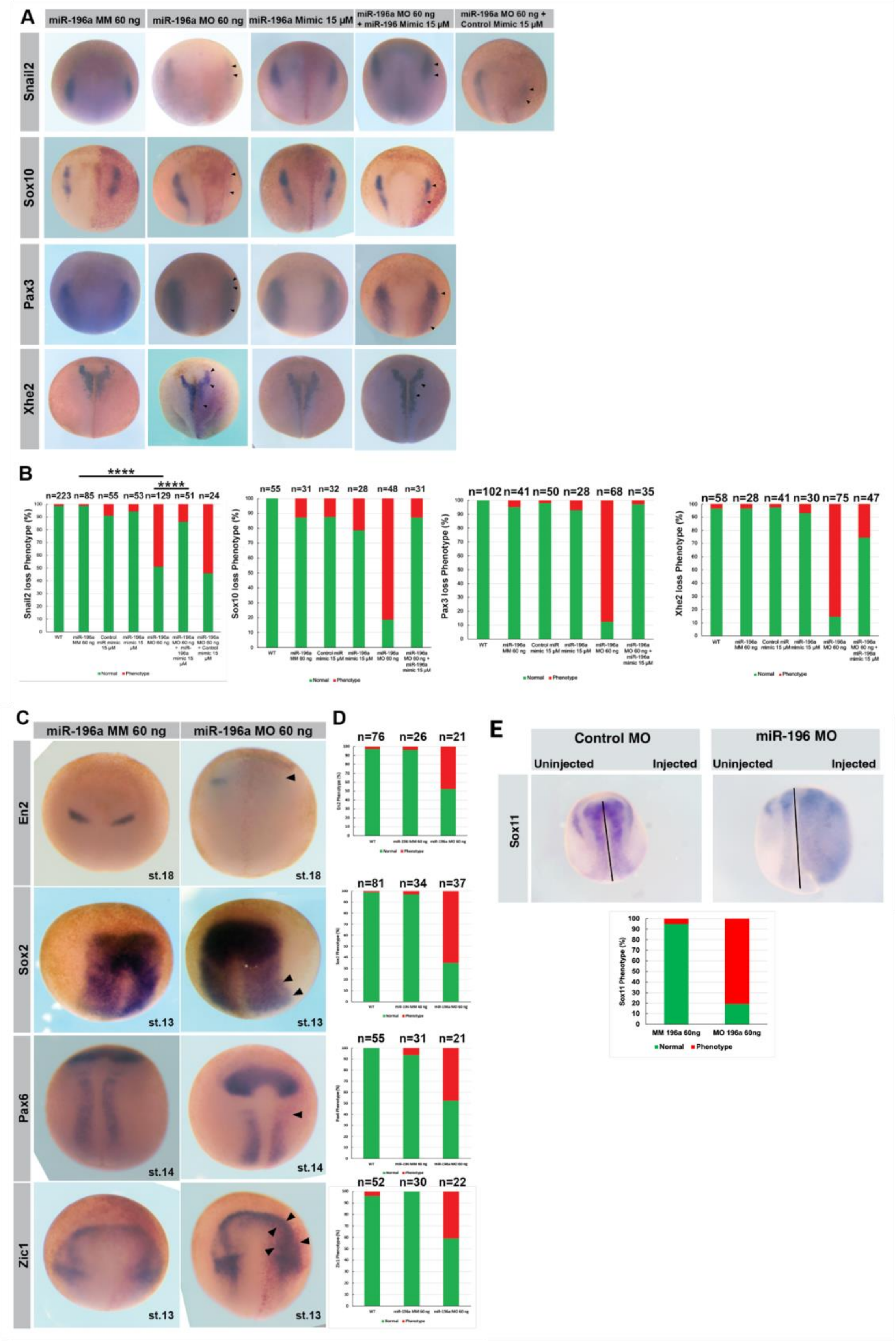

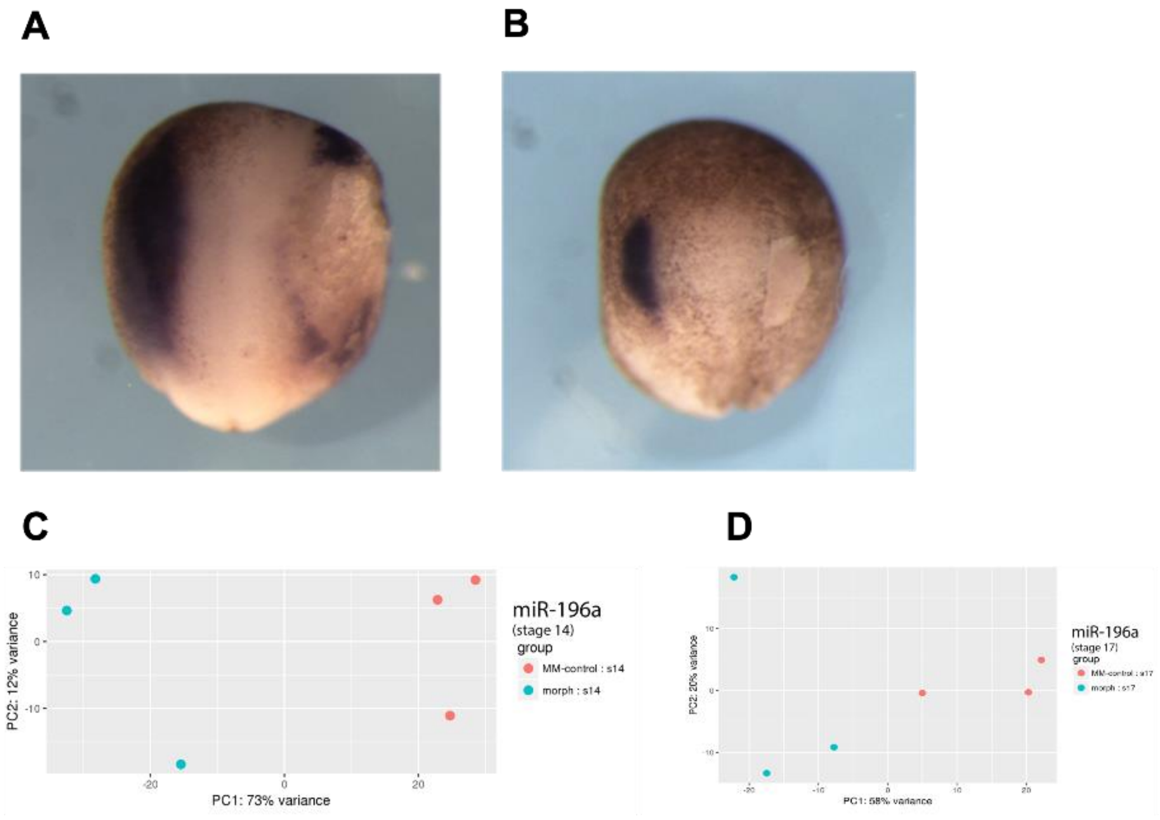

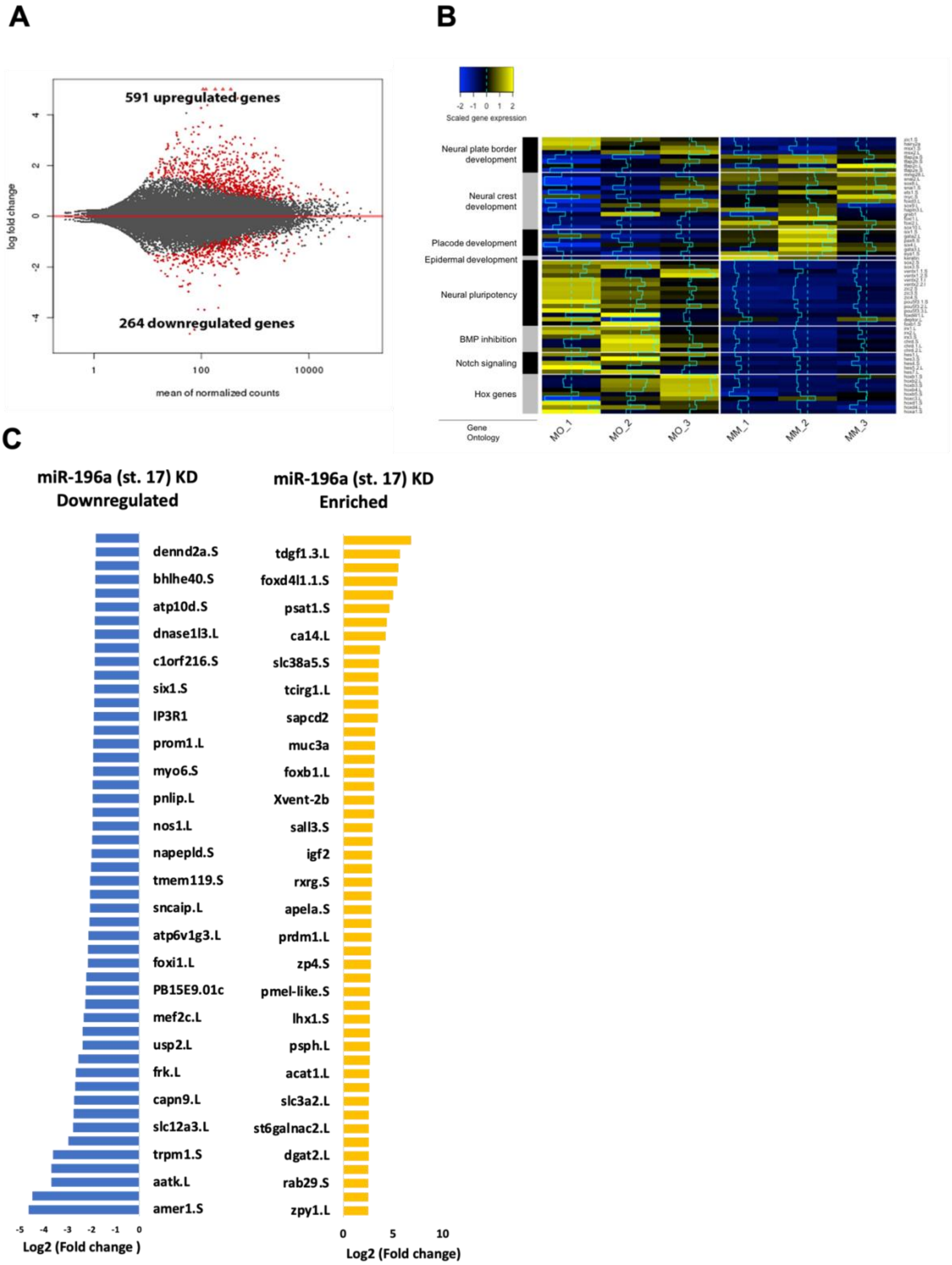
